# PhenoMultiOmics: an enzymatic reaction inferred multi-omics network visualization web server

**DOI:** 10.1101/2024.04.04.588041

**Authors:** Yuying Shi, Botao Xu, Jie Chai, Cheng Wang

**Author notes:** To whom correspondence should be addressed. Jie Chai, Cheng Wang. The authors wish it to be known that, in their opinion, the first two authors should be regarded as Joint First Authors.

## Abstract

Enzymatic reactions play a pivotal role in regulating cellular processes with a high degree of specificity to biological functions. When enzymatic reactions are disrupted by gene, protein, or metabolite dysfunctions in diseases, it becomes crucial to visualize the resulting perturbed enzymatic reaction-induced multi-omics network. Multi-omics network visualization aids in gaining a comprehensive understanding of the functionality and regulatory mechanisms within biological systems. In this study, we designed PhenoMultiOmics, an enzymatic reaction-based multi-omics web server designed to explore the scope of the multi-omics network across various cancer types. We first curated the PhenoMultiOmics Database (PMODB), which enables the retrieval of cancer-gene-protein-metabolite relationships based on the enzymatic reactions. We then developed the MultiOmics network visualization module to depict the interplay between genes, proteins, and metabolites in response to specific cancer-related enzymatic reactions. The biomarker discovery module facilitates functional analysis through differential omic feature expression and pathway enrichment analysis. PhenoMultiOmics has been applied to analyze transcriptomics data of gastric cancer and metabolomics data of lung cancer, providing insights into interrupted enzymatic reactions and the associated multi-omics network. PhenoMultiOmics is freely accessed at https://phenomultiomics.shinyapps.io/cancer/ with a user-friendly and interactive web interface.

## Introduction

Systems biology leverages genomic, proteomic, and metabolomic technologies to unravel the etiology and mechanisms of diseases within intricated biological systems. Integrating diverse biological components, including genes, proteins, and metabolites, enriches our understanding of interconnected cellular activities and the abnormal metabolic states present in diseases (1–4). A fundamental aspect of a cell’s biological omics cascade consists of an extensive set of enzymatic reactions, which are composed of substrates and enzymes, making them the simplest systems to investigate regulatory mechanisms in disease (5–8).

The rapid advancement of omics techniques, such as transcriptomics, proteomics, and metabolomics, enables rapid and precise profiling at the genomic, proteomic, and metabolic levels. Efficiently integrating these molecular components is essential for constructing a comprehensive framework that elucidates the relationship between phenotype and cellular characteristics (9, 10). Notably, computational tools have been emerged to facilitate the integration and visualization of genomic, proteomic, and metabolomic data for mechanistic investigation in cancer, Alzheimer’s disease, and cardiovascular diseases (CVD). For example, Zhou G et al. have developed OmicsNet, a user-friendly web server that allows researchers to effortlessly create and visualize biological networks in a 3D space based on biological features such as genes, proteins, and metabolites (11). MiBioMics was designed to generate correlation maps among multi-omics features and extract relevant variables connecting omics layers to a trait of interest (12). Despite many multi-omics analysis tools have been developed, most of them construct multi-omics graphs based on statistical correlations among genes, proteins, and metabolites. To gain a comprehensive view of the multiomics network, it is imperative to incorporate enzymatic reaction information. Though many enzymatic reaction databases have been developed, the enzymatic reaction based multi-omics network web application is still very rare.

In this study, we designed PhenoMultiOmics, an enzymatic reaction-inferred multi-omics database that integrates 48268 enzymatic reactions across 20 cancer types, using data from the Metabolic Atlas (13). This integration produces a wealth of information, including protein-metabolite associations, 12901 disease-gene associations from the Disgenet database, and 8347 gene-protein associations from the UniProt database. PhenoMultiOmics incorporates a statistical analysis module for conducting differential omic analysis and generating multi-omics networks associated with diseases like gastric cancer and lung cancer. PhenoMultiOmics is hosted using R Shiny web applications at https://phenomultiomics.shinyapps.io/cancer/, offering a user-friendly and interactive interface. As such, PhenoMultiOmics promises to greatly facilitate the study of multi-omics analysis with enzymatic reaction information, furthering our understanding of mechanistic processes in biology and diseases.

## Methods

### Construction of PhenoMultiOmics database

In this section, we outline the process of constructing the PhenoMultiOmics database (PMODB). Initially, we extracted metabolic reactions involving proteins and metabolites from the Metabolic Atlas database (14). Additionally, gene-disease association information was obtained from the Disgenet database (15). Each metabolic reaction in PMODB inherently incorporates gene information, enabling the creation of pairs such as gene-protein, protein-metabolite, and metabolite-disease connections. PMODB comprises essential information about diseases, genes, proteins, and metabolites, each assigned a public database. For genes and diseases, it includes the disease name and ID, gene name, gene symbol, and gene-disease association score. Regarding proteins, the database contains information such as protein name, enzyme classification number (EC number), UniProt ID, enzyme name, protein sequence length, and the corresponding gene name for the protein. As for metabolites, PMODB includes metabolite name, chemical formula, SMILES notation, and associated metabolic pathways.

### Visualization of multi-omics network

Based on the established PhenoMultiOmics Database (PMODB), a multi-omics network is generated utilizing enzymatic reaction data involving genes, proteins, and metabolites. In this network, each gene, protein, or metabolite is represented as a node. The enzymatic reaction information is used to create edges between genes, proteins, and metabolites. Specifically, in enzymatic reactions, edges denote interactions between proteins (enzymes) and metabolites (substrates and products). Additionally, for genes that regulate protein expression, connections are established between genes and proteins, as well as between genes and metabolites. Considering the bidirectional or reversible nature of enzymatic reactions, the network does not contain the direction of reactions, and edge weights are not assigned (16).

### Biomarker discovery module

The PhenoMultiOmics web server incorporates a biomarker discovery module for statistical and functional analysis. For statistical analysis, differential omic feature data analysis is embedded, which require the matrices of gene expression, proteomics, or metabolomics data as input. Each row of this matrix represents a gene or feature, and each column corresponds to a sample ID. This analysis leverages the lima R package to calculate the Log2 Fold Change (Log2FC), estimating differences between case and control groups (17). Student’s t-test is performed to ascertain the statistical significance between these groups, and Benjamini and Hochberg (BH) correction is applied to adjust the P-values. Results are presented in tables and volcano plots, with default criteria for significance being P < 0.05 and an absolute Log2FC ≥ 1 using the provided demo data. Furthermore, for differentially expressed genes, Gene Set Pathway Enrichment Analysis (GSEA) is conducted, drawing from the clusterProfiler package (18). This analysis utilizes human genome annotation from org.Hs.eg.db and pathway enrichment data from the KEGG database (19, 20). Enrichment factors and BH-corrected P-values are computed to identify significant enrichment pathways, displayed via bar and bubble charts. Finally, the enrichplot R package is used to visualize networks of individual genes and pathways.

### RNA-seq transcriptomics data processing

This section describes the dataset used for RNA-seq transcriptomics data, sourced from the publicly accessible database PRJNA152559, GSE35809 (21). The dataset includes genome-wide mRNA expression profiles of 70 primary gastric tumors from an Australian patient cohort, categorized into metabolic (15 samples), invasive (26 samples), and proliferative subtypes (29 samples). Gastric adenocarcinomas exhibit considerable heterogeneity among patients, and the identification of these subtypes can offer insights into cancer progression mechanisms and personalized treatment possibilities.

### Metabolomics data processing

Metabolomics data were extracted from the Metabolomics Workbench (https://www.metabolomicsworkbench.org/) with study ID ST001231. The dataset consists of 31 plasma samples from individuals with lung cancer (experimental group) and 35 plasma samples from individuals without lung cancer (control group). The data were acquired using untargeted metabolomics via UPLC-QE-MS experiments, resulting in a metabolite intensity matrix for all samples, which was used for subsequent statistical analysis.

### Implementation in R

The PhenoMultiOmics web server is programmed in R (version: 4.3.1) and relies on packages provided by Bioconductor and CRAN. The multi-omics network is constructed using the visNetwork package, while the limma package facilitates statistical analysis. GSPEA is conducted using the clusterProfiler package (22–24). All visualizations, including figures and plots presented on the web server, are created with the ggplot2 package and are available for download (17, 18, 25).

### Long-term sustainability of PhenoMultiOmics

To maintain the relevancy and effectiveness of PhenoMultiOmics, we have implemented a version control strategy. Each release of PhenoMultiOmics aligns with specific updates from our key external data sources, Metabolic Atlas and Disgenet. Our approach involves routinely tracking these databases for new data releases and manually integrating this fresh data into our platform. Consequently, every update is meticulously documented and assigned a unique version number. This system allows users to effortlessly identify and select their desired data version, catering to their specific research requirements. Moreover, we commit to ensuring transparency and user clarity by comprehensively documenting each version’s update details, including the nature and timing of data changes, on the PhenoMultiOmics platform. This method not only secures data timeliness but also facilitates user access to a historical data archive for comparative and retrospective analyses.

## Results and Discussion

### Framework of the PhenoMultiOmics web server

The PhenoMultiOmics web server’s framework is illustrated in **Figure 1**, featuring two key modules: PMO database, and PMO biomarker discovery. The PMO database (PMODB) is structured to store data records formatted according to enzymatic reactions involving genes, proteins, metabolites, and phenotypes. PMODB enables rapid access to multi-omics reaction relationship information by querying genes, proteins, or metabolites, with the query results available for download in tabular format. The PMO Biomarker discovery module encompasses differential omics analysis, gene set pathway enrichment analysis, and multi-omics network visualization facilitating the identification of disrupted genes, proteins, or metabolites and enabling functional analysis. Differential expression analysis results are illustrated using volcano plots, while enrichment results are displayed through bubble charts, column diagrams, and enrichment gene network diagrams for each pathway. The PMO multi-omics network, a visualization tool for reaction-based network information, highlights disruptions in genes, proteins, or metabolites related to diseases. In this network, each entity (gene, protein, or metabolite) is represented as a node, with connections based on enzymatic reactions as detailed in the Methods section. **Figure 2** shows a screenshot of the PMO web server, demonstrating the capability to upload node and edge files for custom network diagram creation. Alternatively, users can upload only node files to generate multi-omics networks using the enzymatic reaction database. The interface allows users to customize the network layout and adjust the node size range according to their specific needs.

**Figure 1.**
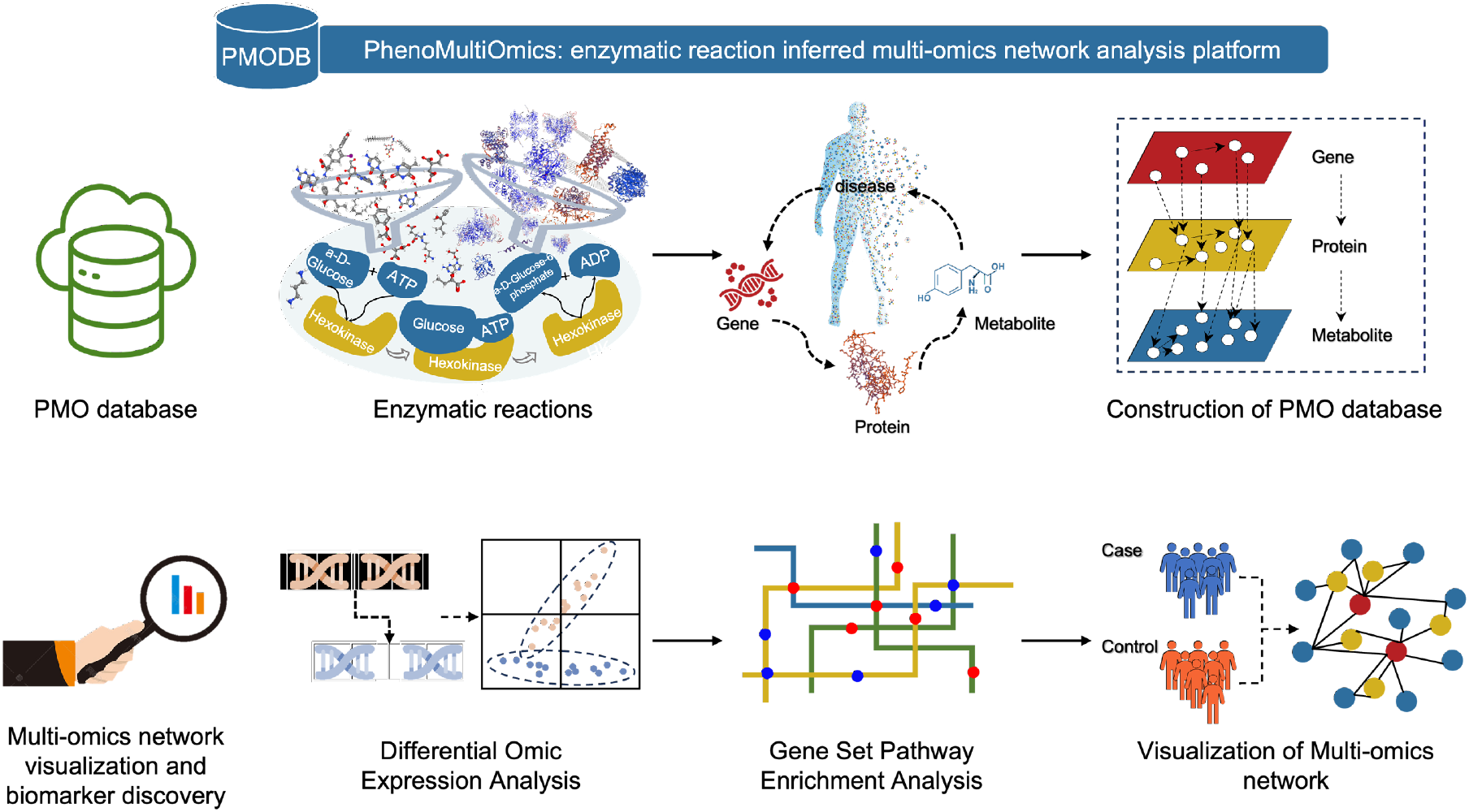
The overview of PhenoMultiOmics database and web server. PhenoMultiOmics constructs a MultiOmics database and network based on thousands of enzymatic reactions. Functional analysis can be performed based on enzymatic reaction data, including Differential Genes Expression Analysis, Gene Set Pathway Enrichment Analysis, and Visualization of multi-omics network.

**Figure 2.**
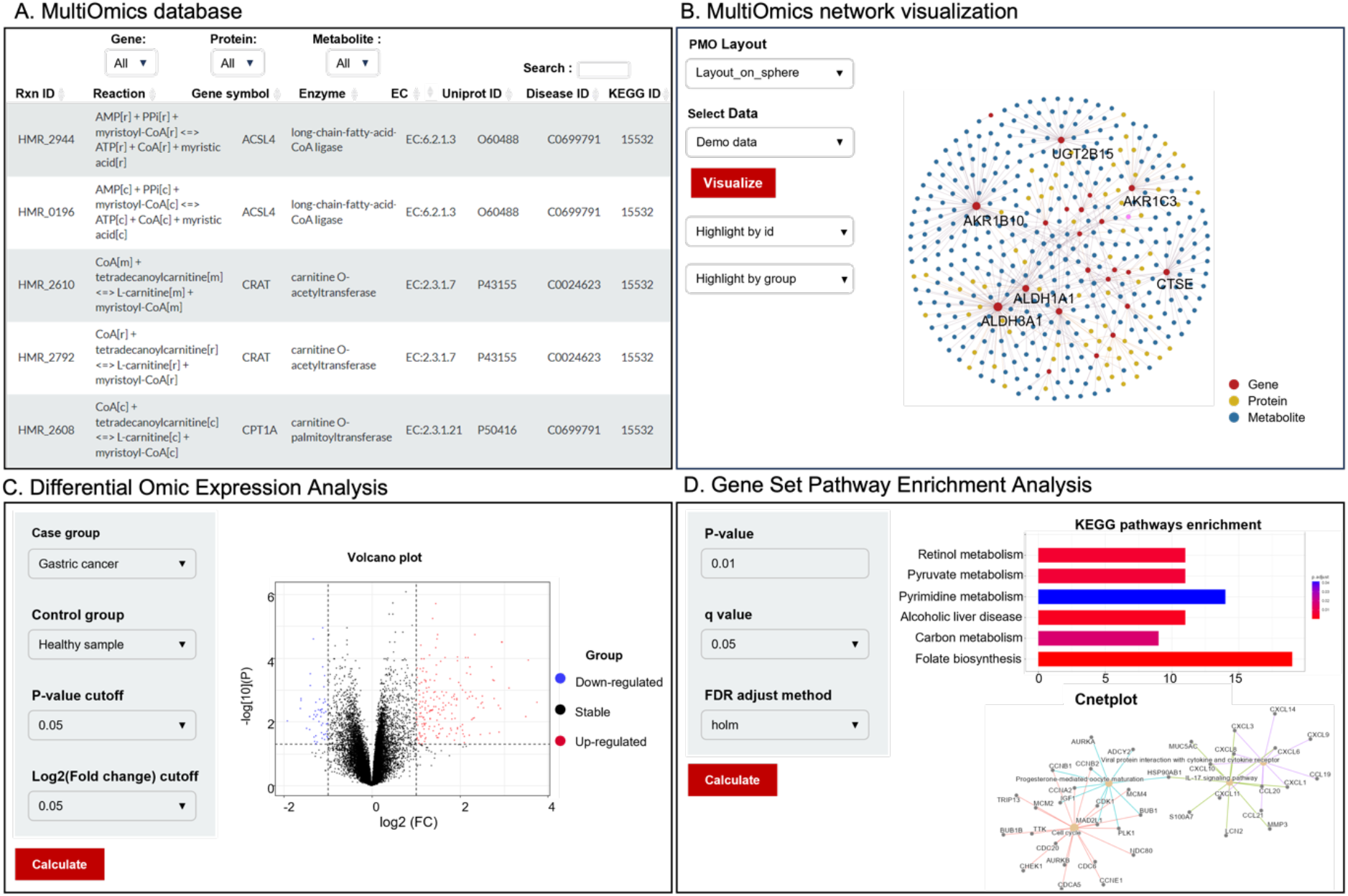
Screenshots of PhenoMultiOmics web server. Panel A shows the MultiOmics database, which enables the retrieval of cancer-gene-protein-metabolite relationships based on the enzymatic reactions. Panel B shows the MultiOmics network to perform visualization of reaction-based network information using disrupted genes, proteins or metabolites in diseases. Panel C shows the Differential Omic Expression Analysis of example data. Panel D shows the Gene Set Pathway Enrichment Analysis using demo data.

### Application to gastric cancer transcriptomics data

Here we applied PhenoMultiOmics to analyze transcriptomics data from different subtypes of gastric cancer, including metabolic, invasive, and proliferative types. **Figure 3** illustrates the outcomes of univariate differential gene expression analysis, revealing significant differences in metabolic enrichment pathways among these distinct gastric cancer subtypes. Proliferative gastric cancer notably associated with gene sets linked to the cell cycle (26). Genome-scale metabolic reaction analysis reveals its enrichment in six pathways: Cell cycle, IL-17 signaling pathway, Viral protein interaction with cytokine and cytokine receptor, Progesterone-mediated oocyte maturation, Gastric acid secretion, and Motor proteins.

**Figure 3.**
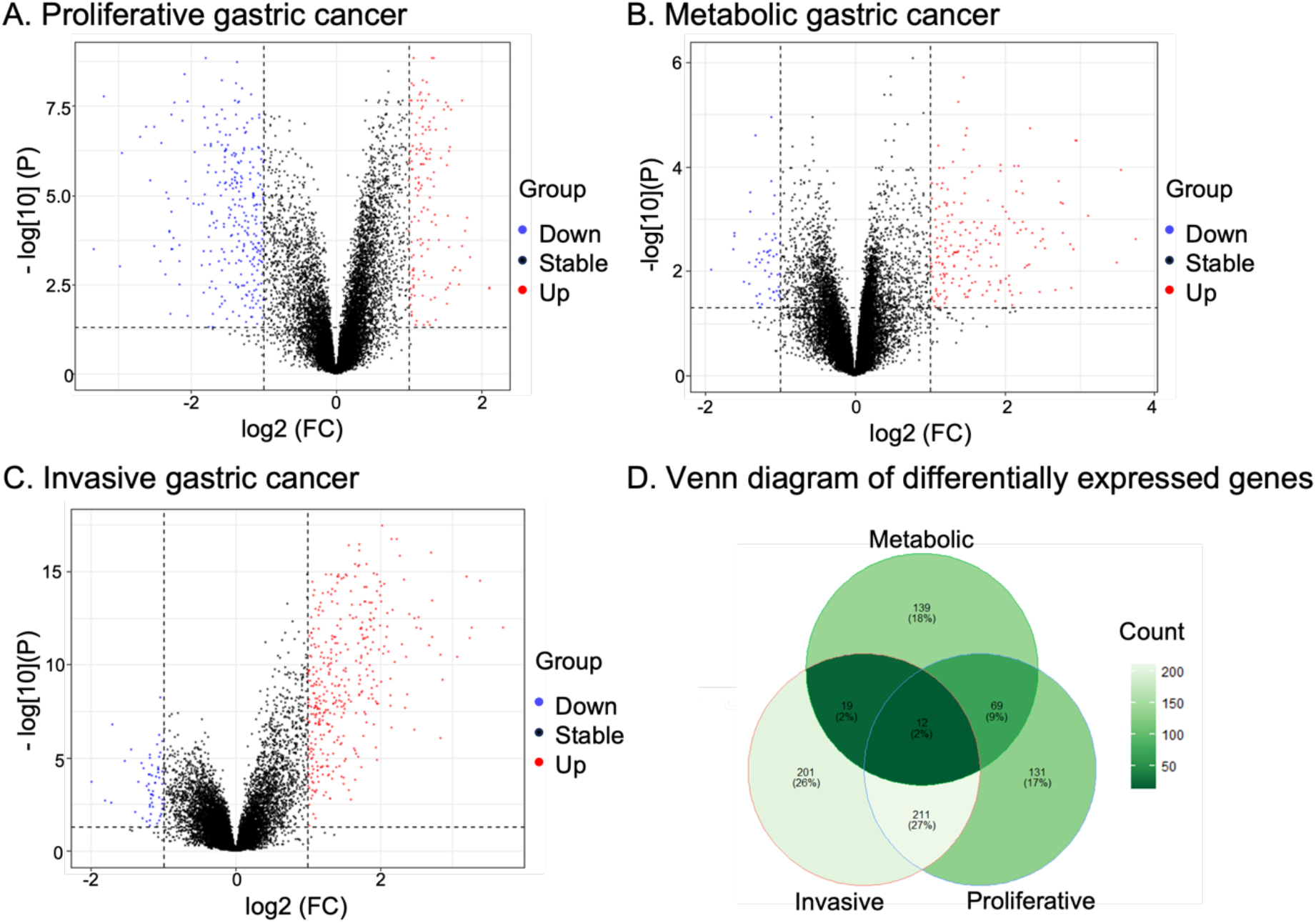
Volcano plots and Venn diagram of differentially expressed gene analysis in gastric cancer. Panel A, B and C show results for different subtype of gastric cancer. Panel D shows the Venn diagram of differential expressed genes in three subtypes, which illustrates the common and distinct genes of three subtypes in gastric cancer.

PhenoMultiOmics multi-omics network analysis identified 2,572 enzymatic reactions, 674 genes, 445 proteins, and 1,941 metabolites related to gastric cancer. Mapping the differential genes of various subtypes onto this multi-omics network provided insights at the gene level for all three gastric cancer subtypes. As shown in **Figure 4**, In these networks, TTK and CDK1 are prominent in modules linked to the Cell cycle and Progesterone-mediated oocyte maturation pathways, both critical for cell proliferation and motility in proliferative gastric cancer (27, 28). Metabolic gastric cancer exhibits gene sets from several KEGG metabolism pathways and GO digestion, particularly enriched in pathways like Retinol metabolism, Metabolism of xenobiotics by cytochrome P450, Gastric acid secretion, Glycolysis, Gluconeogenesis, Fat digestion and absorption, and Vitamin digestion and absorption. Key nodes such as ALDH3A1 and ADH7 in the multi-omics network are notably enriched in Glycolysis/Gluconeogenesis and Metabolism of xenobiotics by cytochrome P450 pathways. Recently, Lei Z et al. identified the sensitivity of invasive gastric cancer to compounds targeting the phosphoinositide 3-kinase-AKT-mTOR pathway (29). Our enzyme-based metabolic network analysis shows the PRKCB gene playing a significant role, enriching pathways like Vascular smooth muscle contraction, Focal adhesion, Gap junction, and the Wnt signaling pathway. Importantly, the Focal adhesion pathway is closely associated with Akt signaling and cell invasion.

**Figure 4.**
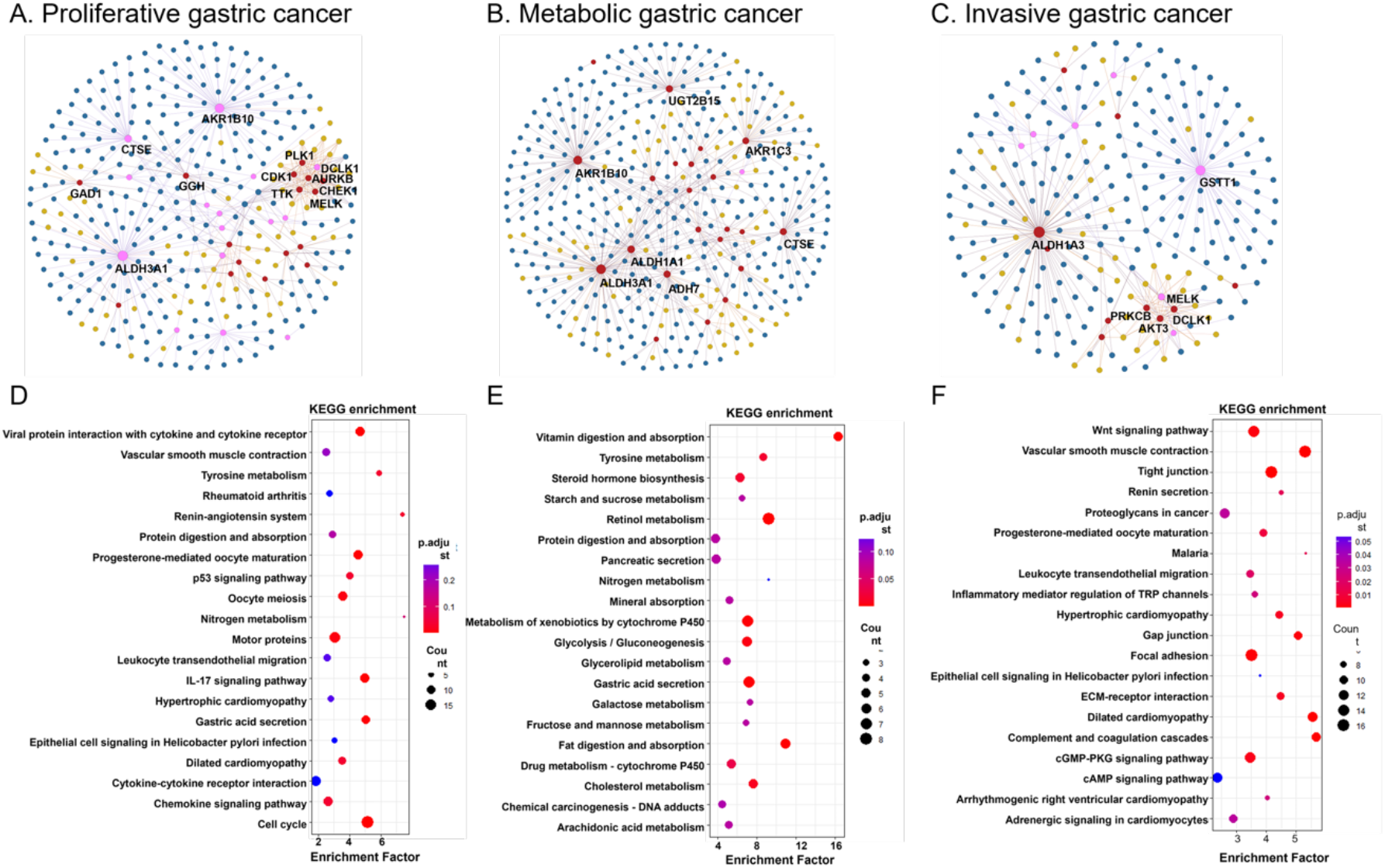
Multi-omics analysis based on differentially expressed genes in distinct subtypes of gastric cancer. Panels A-C show multi-omics network based on enzyme reactions linked to differentially expressed genes in distinct subtypes of gastric cancer. Panels D-F depict enrichment of metabolic pathways of differential genes for three subtypes of gastric cancer. Panels A and D denotes the results of Proliferative gastric cancer. Panels B and E denotes the results of Metabolic gastric cancer. Panels C and F denotes the results of Invasive gastric cancer.

### Application to lung cancer metabolomics data

Next, we extended the application of PhenoMultiOmics to the analysis of metabolomics data related to lung cancer. Our analysis of differential metabolites identified 3940 metabolites. Within the multi-omics database, we identified 185 enzymatic reactions, 42 genes, 49 proteins, and 82 metabolites that constitute the multi-omics networks. Pathway enrichment analysis of these genes, as shown in **Figure 5**, indicated their involvement in 20 distinct pathways. Within the multi-omics network, genes CYP2E1, CYP1B1, CYP1A2, and CYP2A6 exhibit close connections with the enzyme Unspecific monooxygenase. Notably, these genes are enriched in pathways such as Metabolism of xenobiotics by cytochrome P450, Chemical carcinogenesis - DNA adducts, Drug metabolism - cytochrome P450, and Drug metabolism - cytochrome P450. A detailed examination of the Metabolism of xenobiotics by cytochrome P450 pathway revealed the role of cytochrome P450-dependent monooxygenase in seven metabolic reaction pathways, highlighting its potential as a therapeutic target. Inhibiting their activity could disrupt the survival and proliferation of lung cancer cells, thereby enhancing the effectiveness of treatment. Lung cancer onset is closely linked to environmental factors like air pollution, tobacco smoke, and chemical exposures, which often include xenobiotics requiring metabolic processing. Cytochrome P450 enzymes play a pivotal role in handling these xenobiotics, as they convert certain chemicals into metabolites that are more readily excreted. Prolonged exposure to these environmental factors may elevate the risk of developing lung cancer (30, 31). This underscores the scientific validity of extracting multi-omics information related to genes and proteins from the dimension of metabolites through a multi-omics network approach.

**Figure 5.**
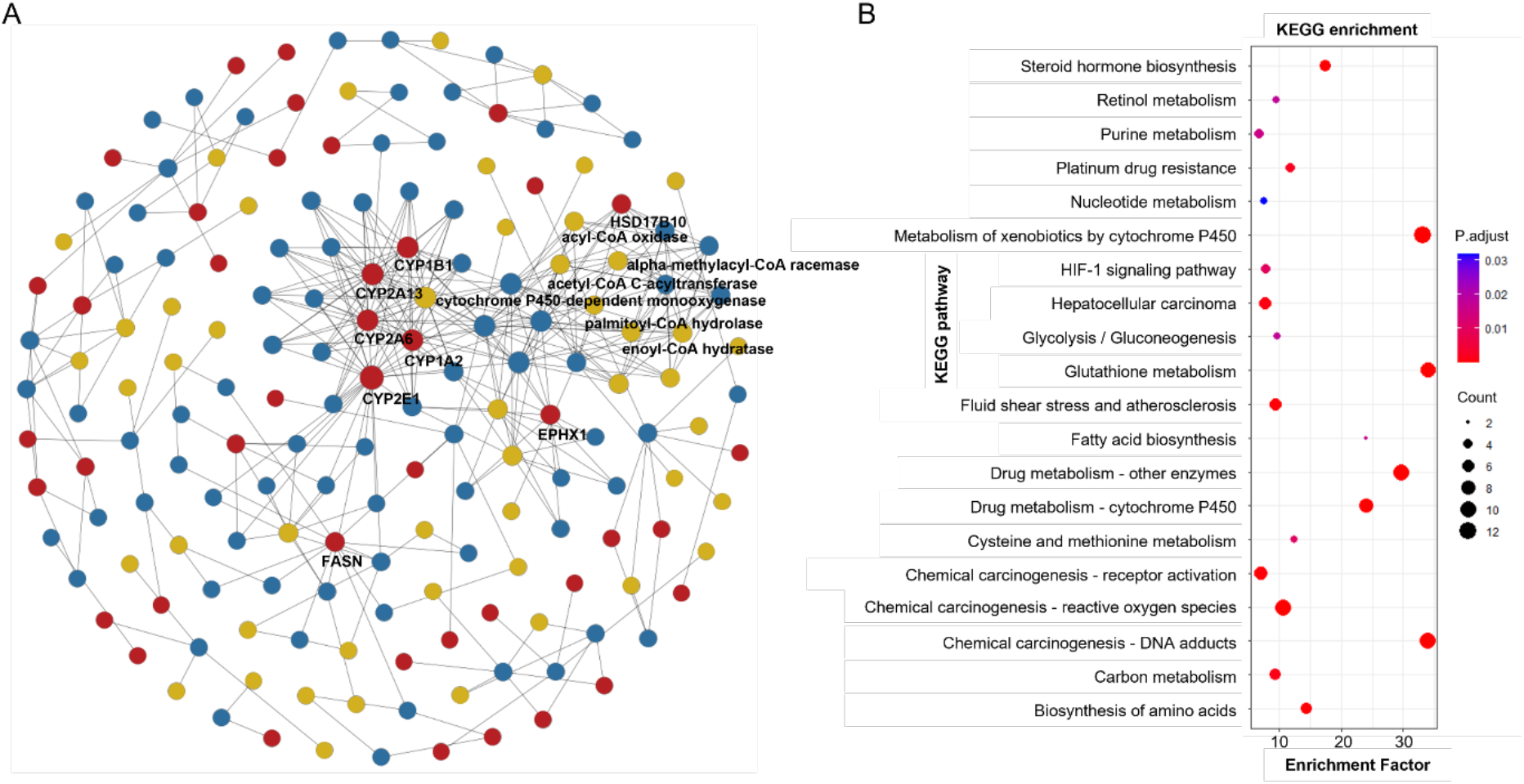
Multi-omics network visualization and functional analysis using disrupted metabolites of lung cancer. Panels A shows results of multi-omics network using the disrupted metabolites based on LC-MS metabolomics data of lung cancer. Panels B shows the results of metabolic pathway enrichment of differentially expressed metabolites in panel A.

## Conclusions

In conclusion, PhenoMultiOmics, a web-based tool anchored in enzymatic reactions, has been developed to integrate statistical and functional analysis for the swift analysis and visualization of multi-omics data. Demonstrating its utility, PhenoMultiOmics was applied to cancer transcriptomics and metabolomics datasets. This tool features a biomarker discovery module enabling differential omics analysis and the generation of disease-specific multi-omics networks, with a focus on gastric and lung cancers. The case studies underscore the tool’s efficacy, showing a high level of concordance between PhenoMultiOmics’ results and the known biological metabolic reactions associated with these diseases. PhenoMultiOmics, by integrating various biological components through enzymatic reactions, is poised to significantly enhance biomarker discovery and deepen our understanding of the intricate cellular activities and altered metabolic states prevalent in diseases.

## Data Availability

The web application is available at https://phenomultiomics.shinyapps.io/cancer/. The source code and data used in the study are freely accessible upon reasonable request to the author (Yuying Shi).

## Acknowledgements

We would like to express our sincere gratitude to the computing resources provided by National Institute of Health Data Science of China.

